# Dynamic instability in nanoscale lipid domains revealed by contact mode high speed AFM: effect of amyloid-β and cholesterol content

**DOI:** 10.1101/2025.01.14.633021

**Authors:** Morgan Robinson, Loren Picco, Oliver D. Payton, Nikolas Zelem, Charlotte Baur, Michael A. Beazely, Mervyn J. Miles, Zoya Leonenko

## Abstract

Cellular membranes are an essential feature of life, the composition and structure of which is important in governing cellular processes and is linked to multiple disorders. Of particular interest is the role that the lipid membrane plays in amyloidogenic diseases such as Alzheimer’s disease (AD), including the role of lipid composition and cholesterol in mediating amyloid toxicity. To mimic neuronal membranes, we used 3-component (DPPC/DOPC/Chol) and 5-component (DPPC/POPC/Chol/sphingomyelin/GM1) model membranes. Atomic force microscopy (AFM) is a key tool in studying the structures of lipid membranes and their interactions with amyloid. Recent advances in contact mode high-speed AFM (HS-AFM) have made it possible to capture dynamic processes at video rate. We used a unique custom-built contact mode HS-AFM to image model lipid membranes and study amyloid-β interactions in liquid. We demonstrate the advantage of using HS-AFM coupled with spatiotemporal variability analysis to capture the dynamic interaction of Aβ 1-42 monomers and oligomers with phase separated lipid bilayers to elucidate the role of nanoscale domains in amyloid-membrane interactions. We show that amyloid oligomer complexes induce greater dynamic instability than monomers, and that low cholesterol membranes are more susceptible to destabilization. Overall, we demonstrate the advantage of HS-AFM to image biological processes on biologically relevant soft samples and discuss tip-sample interactions at high-speed operation in contact mode on lipid membrane models in liquid environment.

**Graphical Abstract:** 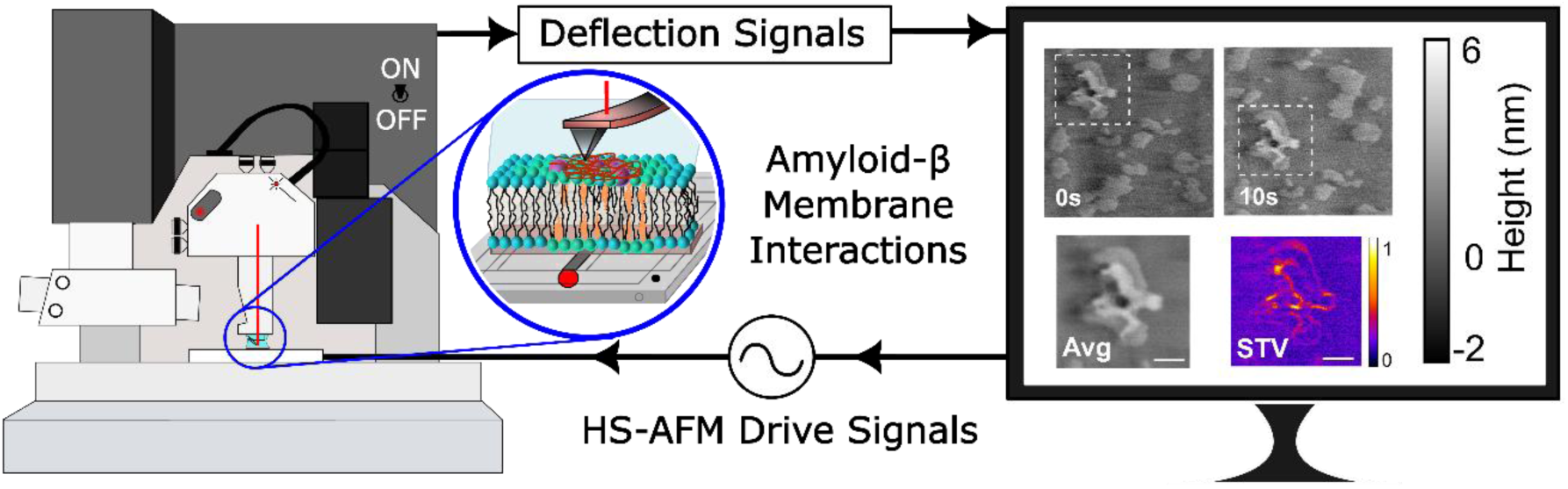

## Introduction

The cell membrane is the critical interface in biology which serves not only to define the extent of the cell, but also, serves to compartmentalize communication and interaction both within the cell and between the cell and the environment. Lipid membrane dysfunction has been implicated in a variety of amyloidogenic diseases, including Alzheimer’s disease (AD), where Amyloid-β (Aβ) has been shown to damage membrane structure which compromises the integrity of the cell, causes toxicity, disrupts ionic homeostasis and interferes with membrane proteins which are important for receptor signaling pathways and neuron signaling ^1–6^. In addition, many physiological processes at the membrane involve non-specific protein-lipid interactions for example: membrane fusion proteins, cell adhesion proteins and binding of amyloid^7–10^. We and others have previously demonstrated that changes in lipid composition directly affect amyloid-membrane interactions and amyloid-induced damage in model lipid membranes, which mimic neuronal cellular membranes in health and AD^11–13^. Thus, strategies to protect the membrane against amyloid damage by modifying membrane properties may be feasible to prevent, slow, or contribute to the reversal of Alzheimer’s disease. This includes protective effects of small membrane active molecules such as melatonin, trehalose disaccharides, as well as modifying the membrane lipid composition^11,14–16^.

Atomic force microscopy (AFM) is an essential tool in nanoscience, especially for analysis of biological samples that require nanoscale spatial precision. However, traditionally it is limited by low imaging rates – on the order of minutes – preventing dynamic biological processes from being imaged^17,18^. AFM generates topographical images by means of mechanical interaction with the surface. Forces between an atomically sharp probe and the sample surface cause a proportional bending of a micrometer-sized cantilever upon which the tip is attached. In contact mode, maintenance of constant tip-sample interaction forces by adjusting the height of the tip relative to the sample as the probe scans over the surface generates a topographical map of the sample. High speed AFM (HS-AFM) systems have been invented that can capture images at video rates visualizing dynamic processes that cannot be imaged with traditional AFM ^19–23^. In this work we used a contact mode HS-AFM developed in Prof. Miles’s laboratory, which not only allows for capturing dynamic changes in topography but also to generate large scale topographical images with extremely high spatial resolution within reasonably short imaging times. This HS-AFM technology has been used for imaging biological samples and capturing processes despite high contact forces with the surface, for instance collagen^24^, DNA in ambient and liquid conditions^25^, as well as the dissolution of tooth enamel from citric acid^26^, and more recently viral fusion on supported lipid bilayers^27^. The contact mode HS-AFM used in this manuscript differs substantially from other high speed AFM systems first pioneered by Ando et al^21,22^, that operate using high-speed tapping mode, and use specialized ultra-short, high resonant frequency (450-650 kHz in water), low spring constant levers. The Ando system tapping-mode HS-AFM technology has been used previously to study amyloid aggregation on mica surfaces^23,28^, amyloid-inhibitor drugs^29^, amyloid-antibody interactions^30^, as well as lipid membrane interactions with carbon nanotubes^31^, amyloid and antimicrobial peptides^32,33^.

In this report, we show that lipid membranes and amyloid-lipid membrane interactions can be resolved at high resolution using contact mode HS-AFM and compare HS-AFM imaging to standard AFM imaging modes. This is noteworthy considering the high contact and shear forces generated by the contact-mode HS-AFM and point to the promise of newer HS-AFM instruments which utilize interferometric optical detection systems for more accurate determination of sub-nanometer height features. We also characterize the interactions between different physiologically relevant Aβ(1-42) species of differing aggregation states (monomeric vs oligomeric) on complex model lipid membranes containing high and low cholesterol. We find that monomeric Aβ(1-42) binds preferentially on liquid ordered domains, generating stable multilayer structures over several hours of incubation. In contrast, preformed Aβ(1-42) oligomers were found to bind less preferentially to the ordered domains, penetrated deeper into the bilayer, forming patches of disrupted membrane, and was prone to solubilizing and removing lipid membrane regions, generating holes in the surface. Aβ (1-42) oligomers also caused greater dynamic instability than monomers, and low cholesterol containing membranes were more susceptible to destabilization as assessed by spatiotemporal variation analysis. These mechanisms of interaction help to explain the relatively greater toxicity that has been associated with Aβ(1-42) oligomers and help to reveal further mechanisms of AD and provide a platform for testing therapeutics for protection of the cell membrane.

## Methods and Materials

### Contact mode standard AFM and contact mode high-speed AFM

A modified version of a previously described HS-AFM system^20,25^ was constructed in-house and mounted onto a Dimension 3100 AFM (Digital Instruments/Veeco/Bruker) which allows for standard and HS-AFM operation, see **Figure 1A** and **1B**. MLCT cantilevers with nominal spring constants of 0.01 N/m purchased from Bruker were used in all cases for both standard and HS-AFM contact mode imaging. The HS-AFM was built using a custom stage scanner with dual sets of flexure bars driven by piezoelectric actuators in both the fast (X) and slow (Y) scan directions to generate motion in the XY plane. The HS-AFM controller (written in-house using LabVIEW) applies a sinusoidal voltage to drive the scan stage in both fast and slow scan axes. Output voltages were sent from LabVIEW through a Test and Measurement Data Acquisition (DAQmx) I/O device (National Instruments) and amplified using a two-channel power amplifier (SONY Electronics Inc.) to drive the piezoelectric actuators of the flexure scan stage. The applied voltages define the scan window size, and the frequency parameters are set according to the desired frame rate. The window scan size was calibrated by moving the Dimension piezo tube a known distance in the X and Y directions while tracking a prominent feature on the AFM movie. The cantilever was mounted onto the Dimension scan tube set to move in a 1 × 1 nm at 1 Hz line rate effectively providing negligible motion in the XY plane while still allowing the scan tube to provide control over the X movement using the Dimension feedback and controller system (NanoScope IIIa or NanoScope IV Controller – Digital Instruments). A constant cantilever deflection was maintained by the Dimension feedback loop set at 1 nN. In-house written data collection software takes the magnitude of the cantilever bending from the Dimension 3100 photodiode via the NanoScope Signal Access Module (Digital Instruments) as the stationary probe sits in contact with the surface via the standard optical lever method – data is then linearized to account for the approximately sinusoidal motion of the scan stage.

**Figure 1.**
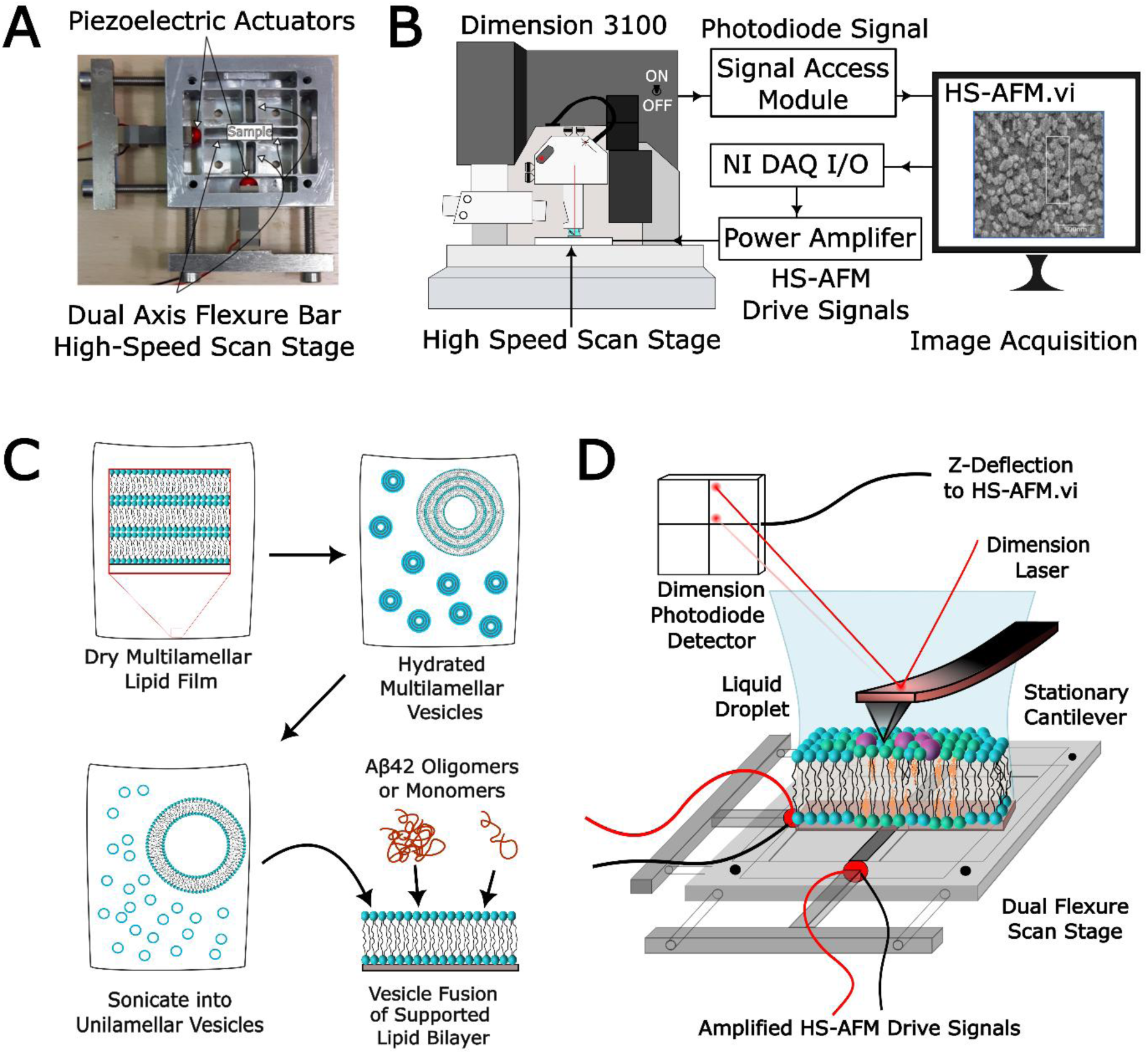
(**A**) HS-AFM dual axis flexure scan stage. (**B**) In-house HS-AFM system integrated with a Dimension 3100 AFM. Sinusoidal AC voltage drive signals are applied to the scan stage piezoelectric actuators from the HS-AFM controller. The amplitude of the AC signal determines the frame/image size, while the frequency governs the image resolution and frame rate. The Dimension 3100 scan tube is set stationary (1nm x 1nm scan window size), and the feedback setpoints are set to maintain surface contact with an average force, below 1nN, cantilever deflection signal is sent to HS-AFM drive and analysis software through a Bruker Breakout Box and plotted in real-time during scanning. (**C**) Supported Lipid Bilayer was produced by sonication and vesicle fusion onto freshly cleaved mica. (**D**) Schematic representation of the lipid bilayer being imaged on the HS-AFM scan stage.

### Lipid membrane composition and preparation

Two model lipid membranes were used in this work: 3-component membrane composed of DPPC/DOPC/Chol at low (45:45:10 by weight) and high cholesterol (40:40:20 by weight) concentration and 5-component membrane composed of DPPC/POPC/Chol/sphingomyelin/GM1 (39:39:10:10:2)^11^. DPPC, 1-palmitoyl-2-oleoyl-sn-glycero-3-phosphocholine (POPC) purchased from Sigma Aldrich, cholesterol purchased from Avanti Polar Lipids, and sphingomyelin and GM1 (monosialotetrahexosylganglioside) sodium salt from bovine brain purchased from Santa Cruz Biotechnology. Supported lipid bilayers were produced by vesicle fusion (Figure 1C), briefly: each lipid was dissolved in chloroform at a concentration of 1 mg/ml mixed at various ratios then evaporated under dry N_2_ gas. Lipid mixtures in chloroform were left overnight to evaporate under vacuum producing a multilamellar thin film of mixed lipid layers. The thin films were then suspended in ultrapure water to 0.5 mg/ml and a SUV solution for vesicle fusion was produced using the sonication method. Vesicle solutions were made one day prior to use, stored in the fridge overnight and then run through 4-8 cycles of 15 min sonication followed by 15 min of stirring immediately, until solutions turned from cloudy to translucent, prior to deposition on freshly cleaved mica by vesicle fusion onto freshly cleaved mica substrates. SUV solution (100 μL) was deposited on mica slide and incubated for 2 hours to establish a stable supported lipid bilayer, before being washed 5 times with MilliQ water (>18.2 MΩ) by exchange 100 µL of solution in the liquid droplet. Supported lipid membranes were imaged in water.

### Monomeric and oligomeric amyloid-β preparation

Aβ(1-42) ultrapure, HFIP treated, (rPeptide) was prepared in monomeric and oligomeric form following protocols previously described^34^. Briefly, dry 1mg Aβ film was suspended in HFIP (Millipore-Sigma) to a concentration of 1mg/mL, aliquots of 0.1mg were prepared by pipetting 100µL into 1.5mL centrifuge tubes, placed in desiccated environment under vacuum overnight. The next day aliquots were placed in -20°C until ready for preparation. Monomeric Aβ was prepared by first making 5 mM solution in DMSO, scraping the sides of the tube and sonicating for 20 minutes in cold water bath. Next, ice-cold MilliQ water was added to the tube and kept on ice. For treatment of supported lipid bilayers, monomeric Aβ was used immediately being injected at 20 µL into the 180 µL liquid droplet for a final concentration of 10 µM. Aβ oligomers were incubated overnight at 4°C overnight, for a minimum of 18 hours, at 100 µM in ultrapure water before being injected into the liquid cell, also at a final concentration of 10 µM. Aβ was injected into the liquid droplet during scanning for some experiments or was incubated on the supported lipid bilayer for 1 hour, before washing 2 times with 100 uL MilliQ water, prior to being imaged. Aβ fibrils were prepared by incubation overnight in 50 µM HCl solution at 37°C overnight as a positive control to validate amyloidogenic capacity.

Preformed Aβ oligomers or monomers were added to the supported lipid bilayer sample, incubated for 60 minutes to allow lipid membrane-amyloid complexes to form, and then excess of amyloid in the bath was washed away prior to imaging in water.

### HS-AFM image acquisition, processing and analysis

For all imaging, the lipid membranes were supported on mica and imaged in water in liquid droplet on the AFM stage (**Figure 1C and 1D**). The standard contact mode images shown herein were acquired using the standard Dimension 3100, with Nanoscope 4 (NanoScope Software version 4). Images were sampled using 512 × 512 px image settings at a line scan rate of 0.5 – 1 Hz, with feedback set below 1 nN of force. Images were then exported into Gwyddion software where they were plane leveled and a Gaussian filter of 3 px was applied. For cross section analysis 5px line width was used. Roughness parameters were calculated by measuring the RMS roughness of each cross section.

For HS-AFM images, in-house LabVIEW Software was used to send drive signals to the high-speed XY scan stage as described earlier. Signals were acquired for processing and sequential display as HS-AFM movies. The frame rate was set to 2 frames per second (fps) using 1,000 Hz fast scan and 1 Hz slow scan drive signals. Electronic phases were manually adjusted to allow for adequate x-y mapping of the Z-deflection signals. The amplitude of the high-speed scan stage (and thus the size of the scan window) was manually adjusted using the gain control of the low impedance power amplifier. Data was linearized to account for approximate sinusoidal motion of the scan stage. HS-AFM videos were then exported from LabVIEW into .avi files and frames were extracted into image sequences using OpenCV packages in python. Individual frames were then imported into Gwyddion and analyzed as above.

For spatiotemporal variation (STV) analysis, image frames were processed using OpenCV pipeline^35^. Frames with amyloid aggregates were identified, the tallest amyloid feature was used to define a template region. This template was then aligned across all frames using normalized cross-correlation–based template matching to identify the amyloid feature across subsequent frames. For each frame, a region centered on the matched location was extracted and saved. This approach enabled consistent tracking and cropping of the same feature across the entire image sequence controlling for drift in the scan region. Only even-numbered frames were used for analysis, as the up (even) and down (odd) frames exhibit hysteresis from the piezoelectric actuators. Next, these cropped amyloid particles were averaged and standard deviation was computed. To compute spatiotemporal variation, which is a scalar measure of the surface topographical dynamics, the average standard deviation (SD) of background lipid surface region and the amyloid particles histograms were plotted. The background SD was subtracted from the amyloid aggregate to account for baseline HS-AFM noise caused by thermal and electrical noise in the cantilever, photodiode sensor, Z- and XY-scanner.

## Results

### Contact mode HS-AFM of supported lipid membranes

The contact-mode HS-AFM system was developed in-house by modification of the Dimension 3100 AFM instrument (**Figure 1A, B**). Calibration of HS-AFM XY scale can be done in two ways; the first is to directly compare identical features under standard and high-speed operation provided a standard AFM image is taken before. Alternatively, under high-speed operation the scan tube can be translated by a set distance across the surface, the resulting translation can be seen and measured by tracking features on the HS-AFM. There are small discrepancies between measuring distances in this way as compared to the XY scale generated from the Dimension under standard operation. Calibration of the Z scale using this modified Dimension 3100 HS-AFM system is less trivial, since the HS-AFM image is generated by plotting the deflection signals directly. The output of the deflection signals from the Dimension piezo tube are calibrated by the Dimension software using a calibration grid, however the native Dimension software does not communicate to the HS-AFM software directly and therefore raw Z-deflections signals cannot be scaled by the HS-AFM software. For the work presented in this report, the HS-AFM system does not have a feedback system with which to calibrate the scale. One way to generate a Z-scale is to compare two identical features captured under standard and high-speed operation and calibrate them. Otherwise force curves on the samples can be taken to calibrate cantilever sensitivity and provide an estimate of the bending angle per unit of voltage on the photodiode detector signal. This limitation of estimating the sample surface heights is not an issue on the latest iteration of contact-mode HS-AFM systems that have recently been developed with laser Doppler vibrometer detection systems which are able to directly monitor the vertical displacement (z height) of the cantilever tip from the sample surface^27,36^.

### Dynamics of 5-component membrane model DPPC/POPC/Chol/SM/GM1

Complex lipid mixtures are often used to mimic native biomembranes and to study the organization and biophysical properties of micro- and nano-domains present in membranes. We used 3 and 5 component lipid bilayers to mimic neuronal membranes in this study. A side-by-side comparison of AFM and HS-AFM images of a complex neuronal model membrane composed of 5 different lipid species was used and shown in **Figure 2A and 2B**; it is composed of: DPPC (36.5% by mole), POPC (35.2 % by mole), cholesterol (17.8 % by mole), sphingomyelin (9.7% by mole) and gangliosides GM1 (0.09% by mole). This complex model lipid system was used previously as biomimetic Alzheimer’s disease (AD) model membrane (Diseased Membrane 1 – DM1) and was shown to be rendered more susceptible to Aβ damage^11^, and be protected from Aβ- induced damage by melatonin^15^. Standard AFM images of one of these model membranes are shown below along with a HS-AFM image of the same membrane and same region in Figure 2. There is a minor x-y distortion of the HS-AFM image when comparing it to the standard AFM due to differences in the exact hysteresis behavior of the piezoelectric stack actuators in the high-speed XY scan stage compared to the XYZ piezoelectric tube scanner in the Dimension 3100 AFM. The difference appears as a compression along the fast scan axis (in the image below the fast scan is in the vertical direction). This error can be corrected using optical interferometric or capacitive sensors to directly measure the trajectory of the scan state, albeit at higher costs^36^. In addition, we observed some minor clockwise skewing of the image which is due to imperfections in the perpendicularity of the slow and fast scan axis piezo, which are not perfectly aligned 90 degrees to one another. Regardless of this minor x-y distortion HS-AFM imaging resolution is comparable to standard AFM. Though there appears to be greater overall image noise, as shown by the cross section in **Figure 2C** and **2D** we found the noise to be variable and dependent on tip and sample quality. Membranes were stable for over 1 hour of continuous recording at 2fps, and exhibited no dynamic changes during that time. Artifacts consistent with scan line mismatch or slow scan drift on standard AFM images were corrected using the remove horizontal scarring feature of Gwyddion, whereas for HS-AFM images, no major line-line artifacts were observed, and horizontal scarring was not corrected.

**Figure 2.**
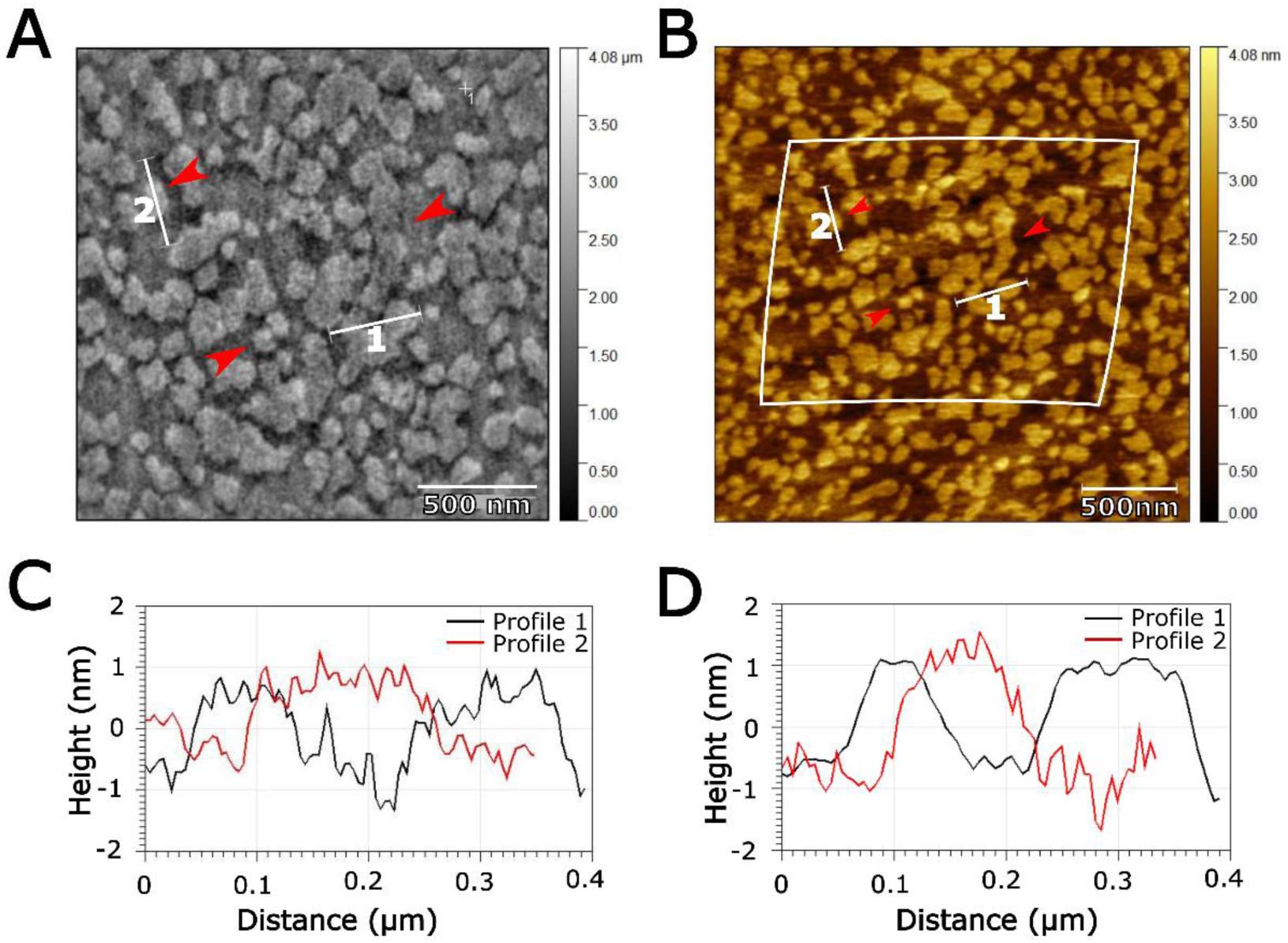
HS-AFM vs standard AFM comparison. (**A**) HS-AFM (2μm×2μm) and (**B**) standard AFM showing the outline of the HS-AFM scan window (2.5μm×2.5μm). Red arrows and cross sections are matched across the two different imaging modalities. HS-AFM scale and vertical axis was calibrated from the standard AFM image extents. (**C**) and (**D**) Cross-section profiles are indicated on the standard and HS-AFM image across L_O_ domains, corresponding to cholesterol, sphingomyelin and ganglioside enriched nanodomains.

### The effect of Aβ (1-42) and cholesterol content on 3-component DOPC/DPPC/Chol membranes

To study the effect of cholesterol content and aggregation state of Aβ(1-42) (monomer and oligomer) on amyloid – membrane interactions, we used simpler, 3-component membrane models (DOPC/DPPC/Chol) at different proportions of cholesterol (40:40:20 and 45:45:10, by weight). Supported lipid bilayers were formed by vesicle fusion, and control images were taken with the HS-AFM system. These lipid bilayers were then treated with 10 µM Aβ (either monomer or oligomer) for 1 hour before being washed 3 times and then imaged again. Sample homogeneity across very large ∼ 2 - 4 mm^2^ areas were determined by manual moving the scan stage with the course adjust, samples with high heterogeneity were discarded, for example: incomplete surface coverage, large holes, or large differences in membrane structure.

For data collection, the HS-AFM scan window size was set to ∼2.5×2.5 µm (500 × 500 px), and frame rate was set to 2fps. After Dimension 3100 guided approach, the scan window was moved in a raster pattern across 10 steps in both x and y directions (for 100 imaging windows). The raster step size was 2 µm, making total area imaged as 22.5×22.5µm, 506.25 µm^2^. At each step, 30-40 frames where captured to collect 15-20s of surface dynamics. Three different regions, 2-3 mm distant from one another, were measured for each sample to capture large scale surface heterogeneity. The total sample area measured for each sample was approximately 1.5 mm^2^, utilizing 3600 imaging frames taken over 30 minutes of recording, for a total of 900,000,000 pixels per sample. There is a trade-off between rapidly mapping large surface areas and capturing dynamic processes at a single location. In this study, we adopted a balanced approach that enabled coverage of a relatively large area while simultaneously resolving short-timescale dynamics at each position.

Representative control DOPC/DPPC/Cholesterol containing lipid bilayers at 40:40:20 (high cholesterol) and 45:45:10 (low cholesterol) by weight (or approximately 1:1:1 and 2:2:1 by molar ratio), are shown in **Figure 3A and 3B** along with their respective cross-sections (**Figure 3C and 3D**). Height profiles between domains were between 0.7 and 1.0 nm for both membranes, similar to what is expected based on previous studies^37,38^. These height differences correspond to the two distinct phases of these membranes, the disordered phase enriched in unsaturated phospholipids, and the ordered phase – enriched in cholesterol and unsaturated lipids. Membranes with lower cholesterol concentration (45:45:10), tended to have more narrow domains that were interconnected across large scan areas, whereas high cholesterol (40:40:20) membranes tended to have more isolated domains. Although this effect was variable, and likely depends on fluctuations in ambient temperature and small variations in lipid ratios from sample to sample, as these models lay very close to the phase boundaries and the melting temperature for the lipids in these samples^37,39^. Variability in lipid domain structure prior to Aβ treatment was observed, likely due to changes in ambient conditions temperature, humidity and small differences in stoichiometry of the lipids, and lipid stability.

**Figure 3.**
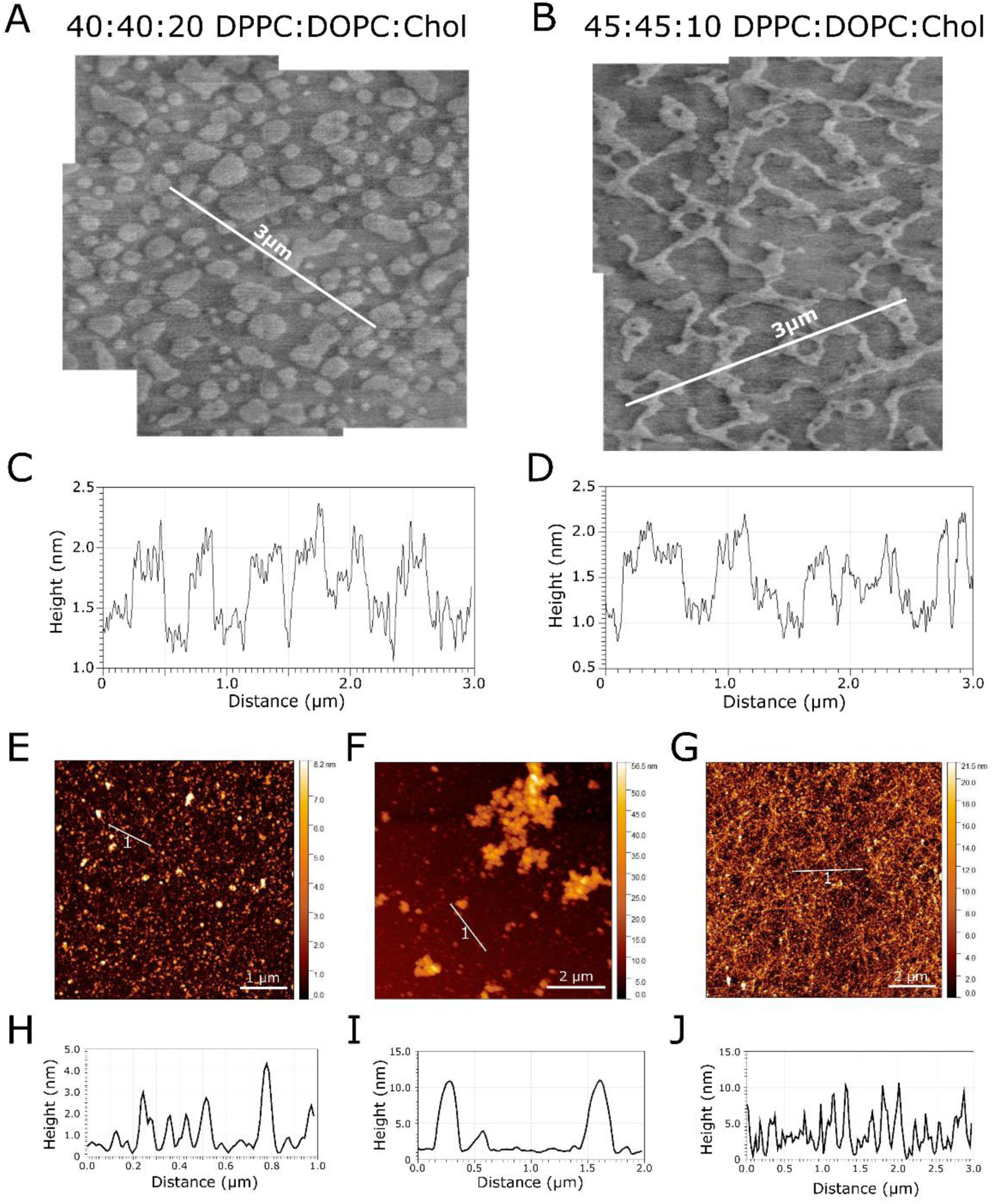
AFM images of DOPC/DPPC/Chol membranes with high and low cholesterol content (40:40:20 and 45:45:10 by weight) and amyloid aggregation states. **(A)** and **(B)** The structure of DOPC/DPPC/Cholesterol lipid bilayers at 40:40:20 and 45:45:10 by weight, with cross sections displayed in **(C)** and **(D). (E)**, **(F)** and **(G)** show representative control images of monomers, oligomers and fibrils on mica substrate, along with the respective cross sections in **(H)**, **(I)**, and **(J).**

After acquiring control images of the membrane models, Aβ as monomers or pre-aggregated oligomers were added to the membranes, incubated for 1h, and then gently washed before the membranes with Aβ were imaged again. Representative monomeric and oligomeric Aβ (1-42) AFM images of monomers and oligomers are shown in **Figure 3E and 3F**, along with a control experiment to show that the Aβ preparations are capable of fibrilization (**Figure 3G**). Cross-sections Aβ show monomers varied in size from <0.5 nm to 4 nm, monomers are likely undergoing some aggregation during their deposition on mica, while oligomers were much larger ranging from 5 - 25 nm in diameter. It is observed that as the HS-AFM scan window moves to a new scan region containing Aβ aggregates on the surface, some of these taller aggregates can be swept off or possibly pressed into the lipid bilayer by the AFM probe. This could be due to differences in binding affinity of the taller aggregates on the surface or due to greater shear forces at larger heights from the AFM probe as it scans over taller image features on the surface. This leaves behind stable aggregates that are more strongly bound to the ordered domain lipid bilayer surface which are stable under high-speed scanning for at least 100 s of HS-AFM frames.

On low concentration cholesterol membranes, Aβ monomers bind preferentially to the ordered taller membrane microdomains, producing structures that protruded above the lower disordered membrane phase by 1.8 ± 0.3 nm after 2 hours (**Figure 4A, D**) and 2.2 ± 0.2 nm after 8 hours (**Figure 4B, E**). Aβ incubation for 2 hours on lipid bilayers produced less tightly bound Aβ aggregates on the surface (**Figure 4A** and **4D**) (N=66 Aβ aggregates across 300 2.5μm×2.5μm frames). After 8 hours, Aβ monomer and lipids appear to organize into very stable multi-layered domains (**Figure 4B**) (N=27 Aβ aggregates across 300 2.5μm×2.5μm frames). At the 2-hour timepoint the RMS line roughness of Aβ on the membrane was increased by 43 ± 18% (N=66 cross sections), then after 8 hours this increased to 70 ± 16% (N=27 cross sections) (**Figure 4G** and **4H**). The increase in surface roughness with time is due to the increased height of Aβ aggregates on the ordered domains in the 8-hour timepoint.

**Figure 4.**
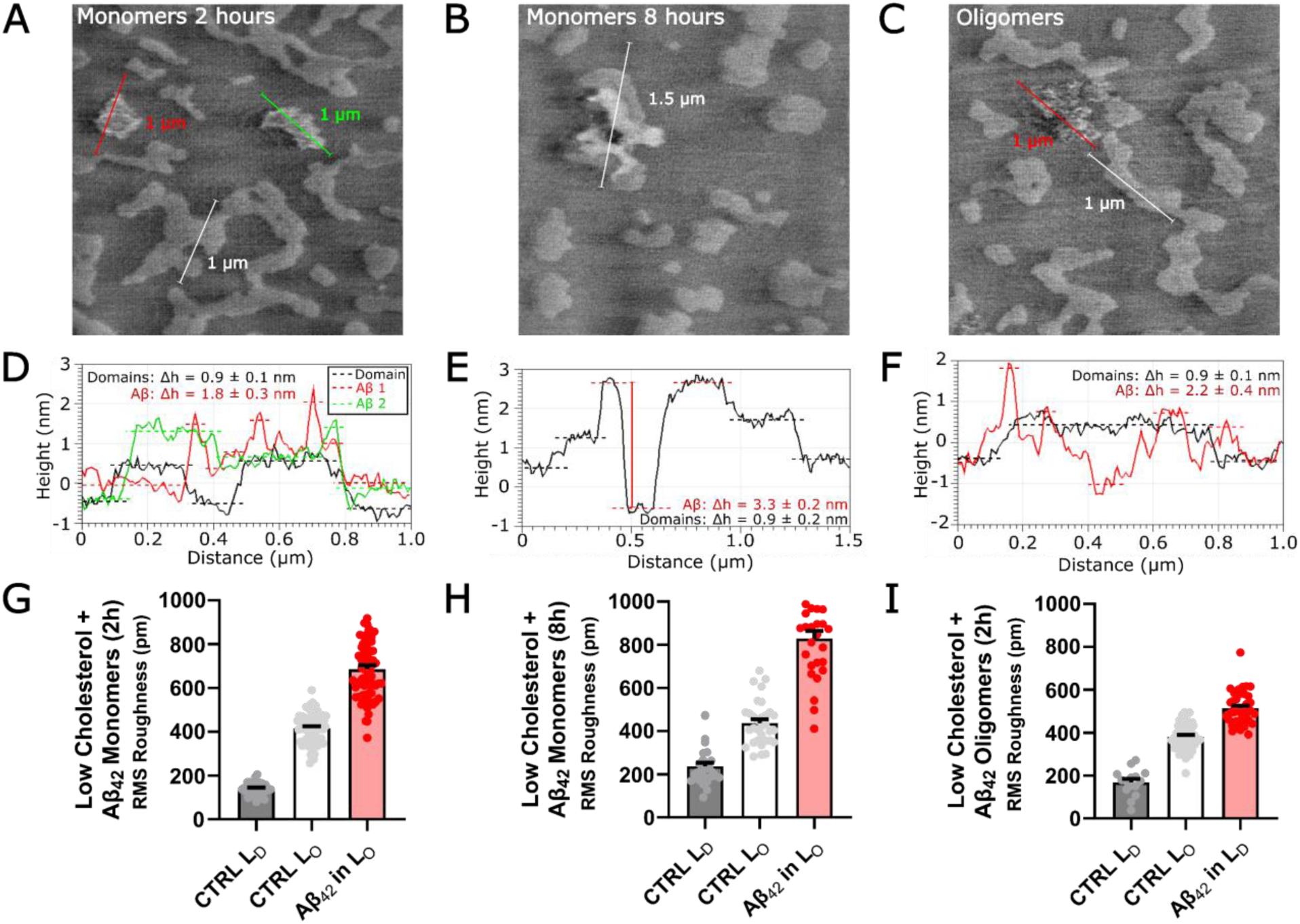
Interaction of Aβ monomers and oligomers with DOPC/DPPC/Cholesterol with low cholesterol content. Low cholesterol lipid bilayers DOPC/DPPC/Cholesterol (45:45:10 by weight) treated with Aβ_42_ for **(A)** 2 hours and **(B)** 8 hours. Aggregates preferentially bind to ordered membrane microdomains and are initially unstable, able to be swept off the surface of the lipid bilayer by HS-AFM imaging after 8 hours aggregates form stable multilayers on ordered domains. **(C)** Low cholesterol lipid bilayer treated with preformed Aβ oligomers. Corresponding Cross-sections are shown below each representative image **(D)**, **(E)** and **(F)**. **(G)**, **(H)** and **(I)** Increase in membrane surface roughness is observed in regions where Aβ monomers and oligomers have bound to and disrupted membrane topography, RMS roughness of the flat Liquid Disordered domains is shown to illustrate background RMS roughness. Each dot represents a single cross section.

Low cholesterol lipid bilayers treated with preformed oligomeric Aβ exhibited unique membrane interactions (**Figure 4C**). Aβ oligomers did not appear to exhibit preferential binding to the ordered membrane phases, Aβ oligomers bound to the disordered and ordered domains indiscriminately resulting in disruption to both phases of the lipid bilayer. In addition, oligomers induced holes alongside the protruding Aβ oligomers in clusters. The height of the tallest oligomers above the disordered phase was 1.5 nm with minimum heights below the plane levelled mean, indicative of holes in the membrane (**Figure 4F**). These aggregates had increased line surface roughness over the ordered domains, by 16 ± 9% (N=38 Aβ aggregates across 300 2.5μm×2.5μm frames), which was the lowest roughness among all the groups (**Figure 4I**) (N=38 cross sections). This is likely due to the aggregates penetrating deeper into the disordered phases of the lipid bilayer, rather than sitting atop the ordered phases as with monomeric Aβ.

We observed several different morphologies of Aβ monomers on high cholesterol containing membranes, again it was observed that Aβ monomers bind to ordered membrane domains, which are most apparent at the boundaries of the domains (**Figure 5A**). We also observe small pore-like structures, which could form as monomers accumulate at the boundary of a small, ordered domain (**Figure 5B**). A more detailed frame-by-frame analysis of the pore-like structure reveals 4 to 5 globular aggregates arranged in a ring, though this was a low probability event (N=2/ out of 29 Aβ aggregates across 300 2.5μm×2.5μm frames). This similar morphology of amyloid ion channels was detected previously, though in that model system the lipid-Aβ preparation was made in proteoliposome deposition, where Aβ was mixed with the lipids in organic solvent, left to evaporate and produce a thin film and then resuspended into a supported lipid bilayer^40^. This proteoliposome system is much less physiologically relevant than the protocol used here, as Aβ is expected to deposit onto cell membranes from the extracellular space. Ordered domain heights are approximately 0.9 ± 0.1 nm above the disordered phase with Aβ monomers protruding to 1.8 ± 0.3 nm above the disordered phase after 2h incubation (**Figure 5C and D**) (N=38 Aβ aggregates across 300 2.5μm×2.5μm frames). Aβ aggregates on the lipid membrane surface increased membrane surface RMS roughness by 63 ± 15% (N=38 Aβ aggregates across 300 2.5μm×2.5μm frames) (**Figure 5I**).

**Figure 5.**
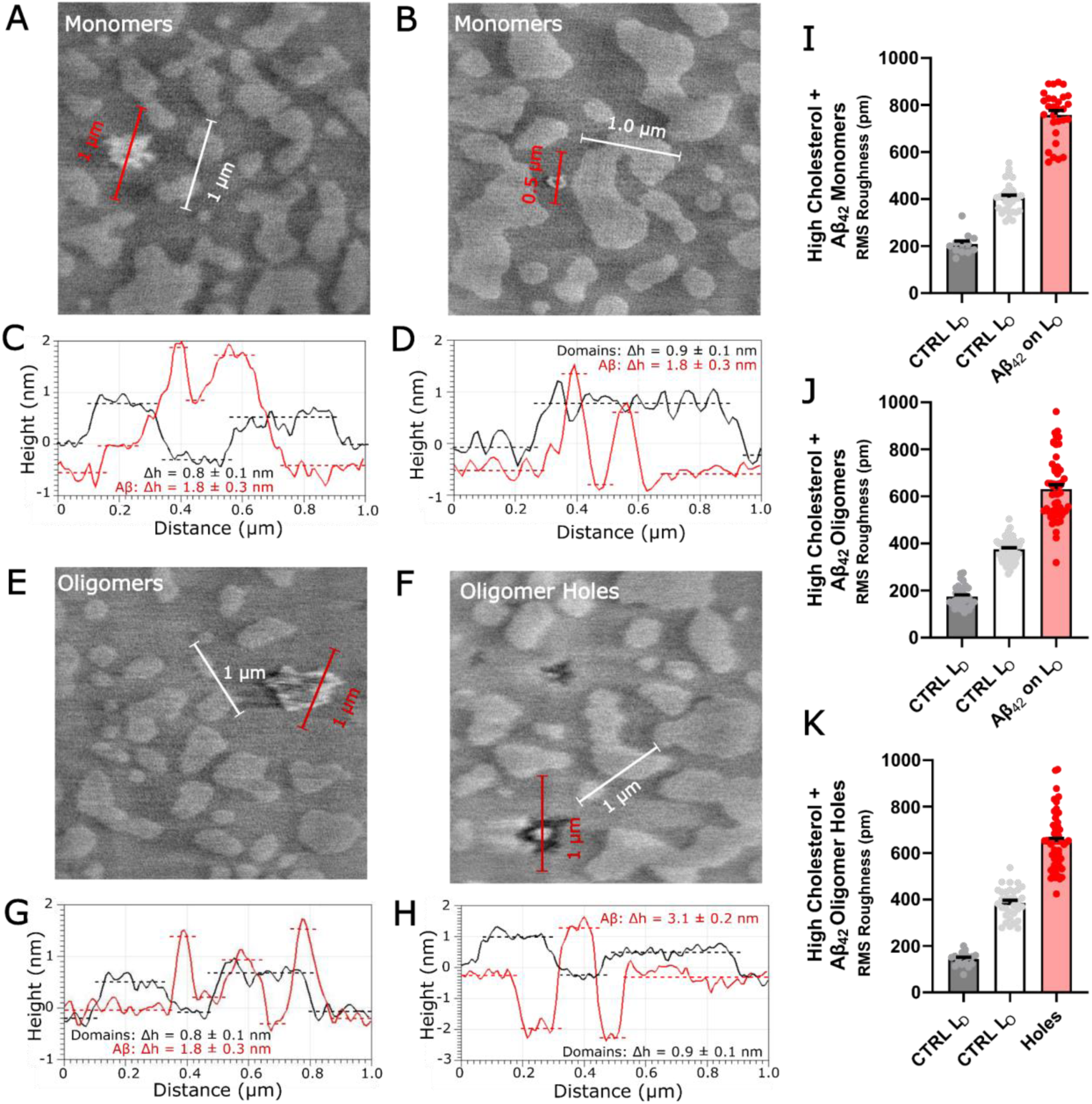
Interaction of Aβ 1-42 monomers and oligomers with DOPC/DPPC/Chol membranes with high cholesterol content. High cholesterol (DOPC/DPPC/Cholesterol (40:40:20)) membranes treated with Aβ Monomers **(A)-(D)** and preformed Aβ oligomers **(E)-(H)**, and roughness analysis **(I)-(K)**. **(A)-(D)** monomeric Aβ was less disruptive to the membrane, forming more stable structures on the ordered domains, cross sections exhibit an increase in domain height over domains not occupied by Aβ. Membrane line RMS roughness for **(I)** High cholesterol bilayer treated with Aβ(1-42) monomers, **(J)** High cholesterol bilayer treated with Aβ(1-42) Oligomers and **(K)** High cholesterol bilayer Aβ(1-42) induced holes; RMS roughness of the flat liquid disordered domains are shown to illustrate background RMS roughness. Each dot represents a cross section from one frame.

High cholesterol lipid bilayers exposed to Aβ oligomers also exhibited more obvious damage to the lipid bilayers in the form of holes more than monomeric Aβ, as in the case of low cholesterol membranes (**Figure 5E** and **5F**). There are two different types of interaction mechanisms observed for Aβ oligomers with high cholesterol membranes. First, binding and aggregation onto the ordered domains like in the case of monomers sitting about 1.8 nm above the membrane surface (**Figure 5G**) (N=55 Aβ aggregates across 300 2.5μm×2.5μm frames). Second, there was also a distinct formation of large and deep holes in the membrane, penetrating as much as 2.1 nm below the ordered phase (**Figure 5H**) (N=55 Aβ-induced holes across 300 2.5μm×2.5μm frames). This type of extraction of the lipids from the membrane surface by amyloid is likely due to some solubilization-like effects that have been reported previously^41^. Typically, bilayer thickness is reported to be at minimum 5 nm, therefore the 2.1 nm depth of the hole could be indicating removal of only the top leaflet or could be a result of tip convolution effects. Regardless, the damage to the membrane is apparent, and distinct compared to all other conditions tested here. Surface roughness analysis does not reveal a difference between the two conditions at 41 ± 17% and 40 ± 2 % RMS surface roughness over the ordered domain to the amyloid containing domain (**Figure 5J** and **5K**) (N=55 Aβ aggregates and N=55 Aβ-induced holes across 300 2.5μm×2.5μm frames).

Low and high cholesterol lipid bilayers treated with Aβ monomers and oligomers exhibit different membrane interactions with some common trends across treatment conditions. In all cases, Aβ increased surface roughness (both mean and RMS), as well as increased height on all the AFM images. Preferential binding of Aβ monomers and oligomers to high cholesterol ordered lipid domains was observed in most cases, except for low cholesterol membranes treated with oligomers. In both low and high cholesterol membranes, Aβ monomers bound to ordered domains and formed aggregates on top of the membrane, in comparison Aβ oligomers were observed to penetrate the membrane and form holes in the lipid bilayer and even remove membrane material.

### Amyloid-induced membrane instability revealed by spatiotemporal variation (STV) analysis of HS-AFM images

Spatiotemporal variation (STV) is a scalar measure of the dynamics of a surface across time. Increased spatiotemporal variation represents larger dynamic instability of the surface. We utilized an OpenCV based computer vision pipeline to extract and align amyloid aggregates, then time averaged, and computed the pixel wise variance of different amyloid species on low and high cholesterol membranes (**Figure 6A**). We analyzed even frames separately from odd frames to ensure that the hysteresis of the piezoelectric actuators did not introduce any apparent stretching/compression in the identified aggregates between the odd and even frames (which is apparent from the contact mode HS-AFM movies in – see **HS-AFM Movie 1**). HS-AFM imaging during raster scanning can also cause stretch distortion which can impair proper blob alignment, and can contribute to apparent motion artefacts, frames with severe motion artefacts were omitted from the analysis. In addition, we observed drift in the piezotube of the Dimension AFM when moving to a new raster scan site. This drift is another well-known problem of piezo actuators, where the tube continues moving in the direction of the voltage, as the voltage slowly decays away over timescales (seconds to minutes), proportional to the magnitude of change in voltage. This drift can affect particle and object alignment, and/or appear as a motion artefact and should be corrected for in the imaging analysis pipeline.

**Figure 6.**
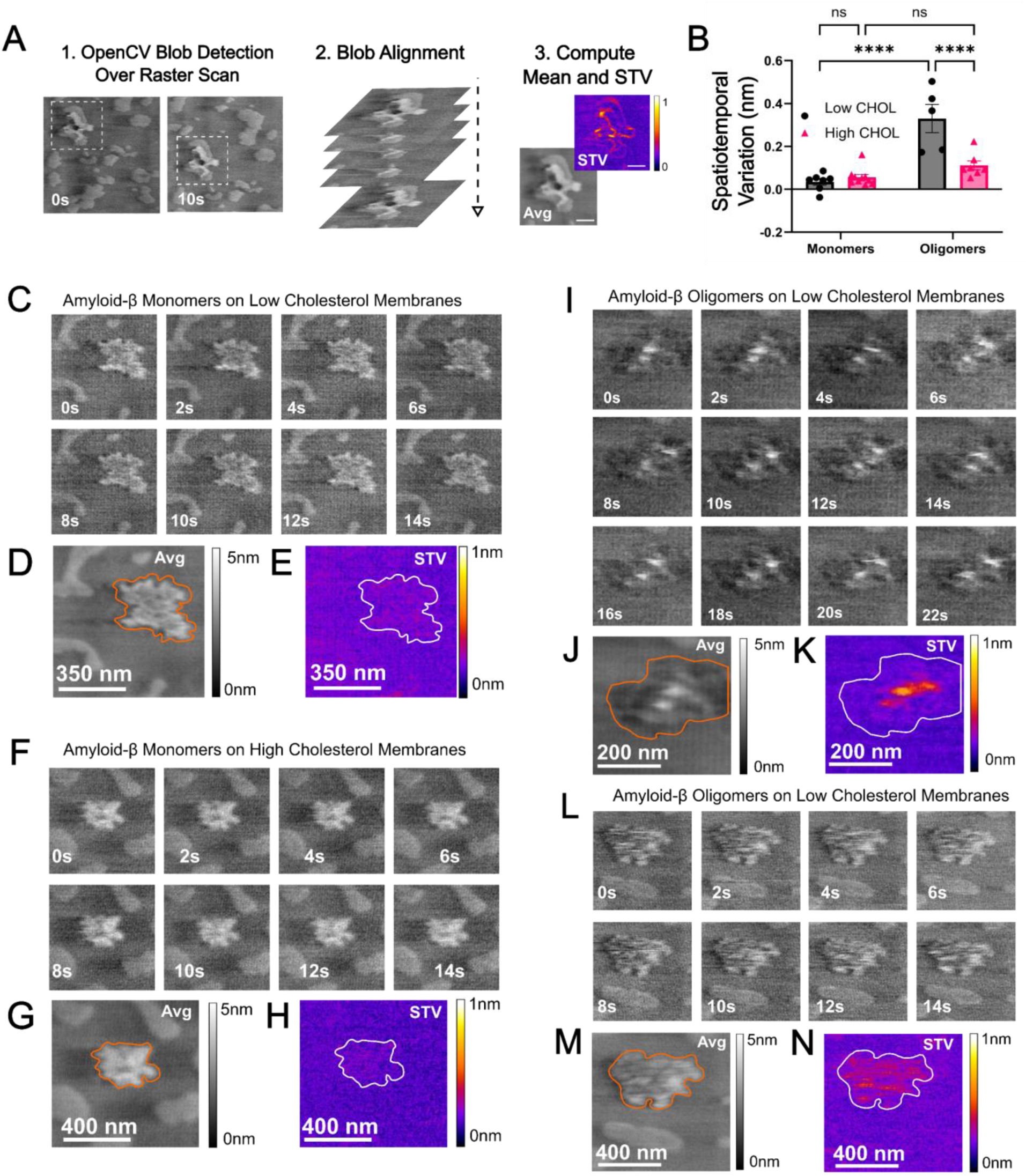
Spatiotemporal variation analysis of Aβ (1-42) monomers and oligomers imaged at 2fps show dynamic instability of lipid bilayers. (**A**) Dynamic image analysis pipeline. (**B**) Overall spatiotemporal variation shows Aβ oligomers cause greater surface dynamics, especially for low cholesterol membranes (two-way ANOVA with Holm-Sidak multiple comparisons). (**C-E**) Representative amyloid monomers on low cholesterol membranes showing even frames along with average image and STV heatmap. (**F-H**) Representative amyloid monomers on high cholesterol domains, time averaged image and STV. (**I-K**) Representative HS-AFM images of Aβ oligomers on low cholesterol membranes, time averaged image and STV. (**L-N**) Representative HS-AFM images of Aβ oligomers on high cholesterol membranes, time averaged image and STV.

Only amyloid particles with at least n=16 frames (16s) were utilized for STV measurements. We found that oligomeric Aβ had higher STV (**Figure 6B**) and thus caused greater dynamic instability of both low and high cholesterol lipid membranes (two-way ANOVA, Holm-Sidak multiple comparisons: monomers N=17, oligomers N=12, ****p<0.001). Interestingly, low cholesterol lipid membranes were more susceptible to destabilization by oligomeric, but not monomeric Aβ (two-way ANOVA Holm-Sidak multiple comparison: low cholesterol oligomers N=5, high cholesterol oligomers N=7), ****p<0.001). There was no statistically significant difference between low and high cholesterol membranes treated with monomeric Aβ.

Representative images are shown in **Figure C-E** (Low cholesterol membranes treated with Aβ monomers), **Figure F-H** (High cholesterol membranes treated with Aβ monomers), **Figure I-K** (Low cholesterol membranes treated with Aβ oligomers – see **HS-AFM Movie 2**), and **Figure L-N** (High cholesterol membranes treated with Aβ oligomers). In the case of Aβ monomers, which were remarkably stable on both low and high cholesterol membranes image averaging was able to resolve amyloid aggregation preferentially on the boundaries of the domains and improved the signal to noise. Most notably, in low cholesterol lipid membranes treated with Aβ oligomers we observed distinct particles which were clearly observable and mobile, contributing to the STV. While high cholesterol membranes treated with oligomers amyloid particles on the ordered domain were more stable than low cholesterol, but less stable than monomeric Aβ.

## Discussion

In this report, we used a custom-built contact mode HS-AFM for studying interaction of Aβ(1-42) with model lipid membranes mimicking neuronal membranes. Typical AFM experiments are long and arduous to collect data, when a single high-resolution image often takes upwards of 20 minutes to gather, then process and analyze. The ability to gather images of comparable quality on the order of seconds means that projects have the potential to be sped up by several orders of magnitude. The contact-mode AFM implemented in this study was able to capture several different interaction modes between different Aβ species, and different cholesterol containing lipid membranes. We showed that Aβ monomers were less disruptive to lipid membranes, preferentially accumulating on ordered domains, forming stable multilayer structures. Interestingly, we observed that Aβ oligomers were prone to causing more damage to both low and high cholesterol membranes, penetrating into the lipid bilayer and extracting membrane material. Spatiotemporal variation analysis further revealed that Aβ aggregates on low cholesterol membranes caused more dynamic instability than in high cholesterol membranes. By utilizing, monomeric and oligomeric Aβ, we were able to mimic different stages of disease progression, with monomers dominating in the early and prodromal AD phases, and oligomeric Aβ increasing during the course of the disease. These results help to better explain the cytotoxicity of Aβ oligomers compared to monomers in cholesterol containing biomimetic lipid bilayers.

Ultra-short cantilever, high frequency tapping mode AFM is another high-speed technology capable of imaging biological membranes that has been used to study Aβ interactions^32,33^. Both the contact-mode work presented here, and previous tapping mode HS-AFM systems are able to capture lipid-protein interactions and study the effects of membrane composition on amyloid interactions. The resolution of each HS-AFM is comparable, though the Ando system may outperform the resolution of the contact-mode system presented here due to the smaller tip diameter^32,33^. However, the contact-mode AFM may have a higher upper limit on the cantilever speed, where we captured 2 fps with scan areas of 2.5×2.5 µm, compared to 2 fps at 800×800nm reported for the tapping mode HS-AFM. Contact-mode HS-AFM lacks phase imaging which can provide additional viscoelastic properties of a sample surface, whereas tapping mode HS-AFM can capture phase contrast images^42^. However, contact mode AFM can be operated in lateral force microscopy or friction force microscopy (LFM or FFM) modes, which are analogous to phase imaging for contact-mode AFM^43^. Friction mapping can be achieved by taking the difference between trace and retrace of the lateral deflection signal, the resulting image then provides a friction map of the surface^43^. Due to the non-linear motion of the high-speed scan stage, sufficiently accurate alignment of the trace/retrace requires direct measurement of scan stage motion using laser interferometry, as has been accomplished elsewhere^36^. Others have mapped friction forces at high speed in contact mode by incrementing the force on the surface and plotting the spatiotemporal difference as a function of the applied force^44^. Overall, the results here show that it is possible to constantly image an extremely soft sample by contact mode HS-AFM to study peptide-lipid interactions with high temporal resolution and larger sample surface area.

Constant contact HS-AFM imaging of soft biological membranes can exhibit destructiveness to poorly fused lipid bilayers. The constant contact high speed scanning was also observed to induce some mixing of the lipid bilayer components under very long imaging times. However, nondestructive high-quality imaging of the lipid bilayer was also shown to be highly reproducible. Non-destructive contact-mode HS-AFM imaging relies on interesting probe-sample dynamics during high-speed scanning: hydrodynamic lift-off of the probe tip and super-lubricity of the ultra-confined water structure between the tip and the sample are proposed as explanations^24^. This HS-AFM is a relatively non-destructive method for imaging soft bilayers with comparable image quality to standard contact AFM mode in liquid. It is possible to limit the effects of the tip over longer-term imaging by taking snapshots and not undergoing constant imaging or scanning over multiple areas spending a short period of time at each location – although these strategies would lower the time resolution of HS-AFM imaging. Newer HS-AFM systems that use laser Doppler vibrometer detection systems and do not have active feedback systems may also be superior for imaging peptide lipid interactions, as the probe height can be kept constant and the force on the surface controlled by height, rather than with a feedback system, which is sensitive to fluctuations in forces during scanning^27^.

Two major advantages of HS-AFM imaging were apparent in our study. First, that HS-AFM imaging allows for much more rapid assessment of sample quality. Within one or two frames of imaging it can be determined whether a complete lipid bilayer is formed, then a rapid assessment of large-scale sample homogeneity over mm^2^ areas can be determined within minutes. This is a process that routinely takes over an hour or longer for standard AFM to image 3-4 areas of a sample to capture the full sample heterogeneity. Secondly, there are less line-to-line mismatch aberrations, and horizontal scarring type AFM artefacts which keep image processing time down. We observed large differences in sample integrity between samples deposited from the same stock solution of lipid vesicles. This points to small differences in ambient conditions, mica surface quality, and lipid stability in solution as difficult to control factors in supported lipid bilayer quality.

Alzheimer’s disease is a chronic condition characterized by slow accumulation of amyloid over the course of decades^45–47^. Previously, tapping mode HS-AFM was used to study the initial binding amyloid-β interactions with biomimetic membranes composed of SM/POPC, SM/POPC/Chol and SM/POPC/Chol/GM1 interact with amyloid. The experimental design of those reports added Aβ to the membrane surface during imaging mimicking the acute toxicity of Aβ oligomers^32,33^. They showed that the inclusion of GM1 was necessary for Aβ-G37C oligomer binding, and that mutant Aβ-G37C peptide oligomers caused rapid dissolution of the membrane shortly after addition only in the presence of both GM1 and cholesterol. A follow up study evaluated various Aβ mutations and amyloid species and analyzed their interactions with their model lipid membranes containing GM1/POPC/SM/cholesterol^32,33^. In a complementary approach to these studies, we used contact mode HS-AFM to determine how different forms of amyloid-β monomers and oligomers, as well as varying levels of cholesterol, influence dynamic processes after accumulation. This mimics two phases of Alzheimer’s disease progression, the early stage where monomeric amyloid dominates and the later stage where oligomers are more abundant. Cholesterol metabolism has been implicated in AD through metabolic and genetic risk factors (most notably ApoE4 alleles)^45^, though many uncertainties remain surrounding the role of cholesterol in AD pathogenesis. The cholesterol dependence of Aβ toxicity has been evaluated in several studies showing opposing effects in different model systems, in some studies cholesterol depletion is shown to be protective^2,6^, while others show that cholesterol enrichment in membrane domains is protective^48^. These discrepancies could be due to differences in the aggregation state of amyloid or the lipid membrane composition of different cell types. Our study here points to dynamic membrane processes as a contributing factor in the toxicity of amyloid. We show that monomeric Aβ caused lower dynamic instability to the lipid membrane, compared to oligomeric Aβ which in general exhibited greater dynamic instability. This difference in toxicity could be due to the differences observed in this HS-AFM study, that oligomers are more likely to penetrate and form holes in the membrane, whereas monomers can bind to the membrane surface but are not able to damage the membrane.

This work extends the applications of contact mode HS-AFM to very soft lipid bilayer samples for studying protein aggregation and exploring an important question in molecular neurodegeneration. This work contributes to the understanding of different interaction mechanisms that can occur between different amyloid-β species and lipid membranes of differing cholesterol concentration. The role of membrane composition in the cytotoxic mechanisms of amyloids in general appears to be an important factor^49^. There were large differences in the interaction mechanisms, as indicated by the differences in surface topography, between the different cholesterol levels and the aggregation state of Aβ. These differences are time-dependent and thus rates of molecular diffusion and proteo-lipid organization are likely important in Aβ toxicity.

## Acknowledgement

We acknowledge the International Strategic Partnership Grant from the University of Waterloo (awarded to Zoya Leonenko) and the Collaborative Initiative Grant from the University of Bristol (awarded to Mervyn Miles) for funding this work, as well as MITACS Globalink, the Ontario Graduate Scholarship and Waterloo Institute of Nanotechnology Nanofellowship awarded to Morgan Robinson. As well as Natural Science and Engineering Council of Canada (NSERC) operating grants awarded to Michael Beazely and Zoya Leonenko, and NSERC undergraduate research award (NSERC-USRA) awarded to Nikolas Zelem. We would also like to acknowledge Dr. Liz Drolle for discussions on complex lipid membrane preparation, Dr. Robert Richardson and Dr. Annela Seddon for use of their laboratory prep space and Mr. Ravi Sharma for assistance with data collection.

## Contributions

Conceptualization and project design by Zoya Leonenko and Mervyn Miles. HS-AFM design and build by Loren Picco, Oliver Payton and Mervyn Miles. Lipid and protein protocols by Morgan Robinson, Michael A. Beazely and Zoya Leonenko; Imaging and image analysis by Morgan Robinson and Mervyn Miles; manuscript written by Morgan Robinson with contributions from all authors. Spatiotemporal variation (STV) analysis of HS-AFM images was developed by Morgan Robinson, with assistance from Nikolas Zelem and Charlotte Baur, and supervised by Zoya Leonenko.

## Conflict of Interest

The authors report no competing conflicts of interest related to this work.

## Data Availability

The data supporting this article are included in the main text or have been included as part of the supplementary information.

## Notes

### Competing Interest Statement

The authors have declared no competing interest.

### Summary of Updates

Additional spatiotemporal variation dynamics results is presented in the new Figure 6, along with additional discussion of these results. Minor spelling, grammar corrections have also been made.

